# PanvaR: An R package for fine-mapping and visualizing results from genome-wide association studies

**DOI:** 10.64898/2026.07.15.738769

**Authors:** Collin Luebbert, Rijan Dhakal, Phillip Ozersky, Scott Lee, Todd Mockler, Ivan Baxter

## Abstract

Genome-wide association studies (GWAS) use statistical models to correlate single nucleotide polymorphisms (SNPs) to a phenotype of interest. This scan of the entire genome identifies regions of association with a phenotype, but due to linkage disequilibrium (LD), GWAS on their own cannot identify single genes responsible for phenotypic variation. Rather, fine-mapping of GWAS regions is required, necessitating the use of additional tools and software. With the introduction of more pangenomic resources in a number of crops (Guo et al. 2025; Hufford et al. 2021), the fidelity of these fine-mapping efforts is growing, presenting the opportunity to leverage new information about allelic variation towards gene discovery (Shi et al. 2023; Della Coletta et al. 2021). Panvar is a tool developed to integrate existing software and resources to perform GWAS and fine-mapping in one seamless step. For each identified GWAS peak, panvaR outputs information about LD and SNP effect prediction for each SNP and by layering locations of nearby genes, creates a refined list of possible candidate genes. We have implemented Panvar as an R package, “panvaR”, which runs the analysis functions, creates interactive and static visualizations, and outputs results tables. This tool seeks to bridge the gap between GWAS and gene speeding up an important step of quantitative genetic studies.

## Introduction

Genome-Wide Association Studies (GWAS) have proved powerful for identifying regions associated with a wide range of quantitative traits in plants (Tibbs Cortes, Zhang, and Yu 2021) and the resulting marker associations are a key resource underlying the success of marker-assisted selection in plant breeding (Collard and Mackill 2008). However, as the adoption of gene editing technology grows, the identification of specific genes is becoming increasingly important for crop improvement (Voytas and Gao 2014). Additionally, with genomic sequencing becoming ubiquitous, researchers are increasingly able to leverage insights from synteny and evolutionary conservation to translate functional gene assignments across species (Whitt et al. 2023).

GWAS rapidly screen an entire genome for regions that associate with a phenotype of interest, but because of linkage disequilibrium (LD), neighboring single nucleotide polymorphisms (SNPs) are correlated, leading to Manhattan plots’ characteristic “peaks”. Depending on the species and the size of LD blocks, a peak may contain up to hundreds of genes. Consequently, fine-mapping to prioritize the most likely causal gene for the putative association is essential to identify a causal gene underlying a GWAS peak.

Recent advances in sequencing have led to the creation of many pan-genome datasets, where all lines are fully sequenced and the near-complete inventory of polymorphisms can be known and evaluated for phenotypic consequences (Morris et al. 2026) (Hufford et al. 2021) (Guo et al. 2025) (Kang et al. 2023). However, pan-genome datasets usually include thousands of polymorphisms within each GWAS peak and are difficult to sort through (Tao et al. 2021). Integrating the different types of data that are needed for this fine mapping step using dense marker sets is a non-trivial task and has not been implemented into a species-agnostic visualization tool.

The tool described here, “Panvar”, seeks to overcome this data integration hurdle, speeding up fine-mapping and paring down the list of possible candidate genes underlying a given GWAS peak.Within the Panvar workflow, multiple data types are layered, and quantitative and qualitative filtering steps are applied to extend GWAS results. The flexibility of the package is intended to allow quick integration and visualization of arbitrary, user-defined metrics of SNP and gene importance. Exploratory analysis and subsequent verification have demonstrated Panvar’s capacity to capture expression-level signals, which can help refine GWAS results to identify potential candidate genes. Our test cases show that Panvar can produce successful outcomes across different species and phenotypes.

## Materials & Methods

### Workflow

PanvaR works on a single locus at a time and integrates data around this position to display information at the SNP and gene levels. Required inputs for the program are: 1) a SNP file in vcf or plink format, 2) a gene annotation file that gives locations and annotations of genes, and 3) a specific locus of interest, usually a SNP. The workflow then proceeds by first formatting inputs and optionally re-running GWAS, then generating tables at the SNP and gene level around the QTL and finally, plotting these results. The program relies on two main dependencies: plink2 (Chang et al. 2015) for genotype file manipulation and LD calculation and rMVP for GWAS functionality (Yin et al. 2021).

### Formatting inputs

The first step of the program is to generate standardized inputs for downstream functions (*make_panvar_inputs()*). This uses plink2 to apply optional quality control filters to the genotype file. Users can select a minor allele frequency and missing rate threshold. This also standardizes column names and input files for GWAS using an rMVP input standardization function.

### Running GWAS

While panvaR is meant to be run after GWAS, there are still many common scenarios where it may be desirable to re-run GWAS. This is explored further in the discussion. rMVP is used as the GWAS implementation and generalized linear model (GLM) and mixed-linear model (MLM) outputs are supported (*panvar_mvp_gwas()*).

### The tag SNP

An important step of fine-mapping is determining regions of interest or QTL. Typically this is done by grouping peaks by a combination of p-value, LD and distance to create groups of SNPs that represent discrete loci displaying association with the phenotype (*snp_make_clumps()*). A function is presented to help the user group their output into loci which can be analyzed by panvaR one by one. A single representative SNP, referred to in the package as the “tag snp”, centers the locus and is usually the top p-value SNP in an identified locus.

### Generating tables

PanvaR allows the integration of different types of data at this point, both qualitative and quantitative and at the SNP and gene level. Measures of impactfulness at these two levels can be added to the GWAS results and annotation table to be carried through the analysis. A major functionality of the package at this step is to aggregate snp-level measures of impact or importance to the gene level. The program will collapse SNPs within a gene start and end, and plus or minus an optional buffer of physical distance, into a single value per gene (**supplemental figure 1**). This gives a more compact summary of the maximum impactful measure for a given gene (*make_panvar_tables()*).

### SNP scores

At the table generation step, panvaR can also generate scores at the SNP level which can then be aggregated to a gene level to summarize an arbitrary number of quantitative impactfulness measurements. To generate the score, metrics are first oriented in the proper direction. The final score will be interpreted so that a larger value indicates a more impactful SNP. Metrics where smaller is better (for example p-value) need to be reversed. Next, a min-max normalization is applied to scale metrics to (0,1). Then the mean is taken to generate the final result. An option is supplied to weight this mean by each metric if a user wants to prioritize certain measurements. By default, if a user requests scores to be computed, the program uses p-value, distance to the tag SNP, and LD.

### Plotting

The final output of the program is a plot which provides an information-dense visualization of the tables generated in the previous step (*plot_panvar()*). This flips the traditional manhattan plot 90 degrees and arrays genes located in this physical range alongside. Colors of points in the manhattan plot can be assigned to a user-defined variable and a variable can be assigned to the shape of the points as well. A rug plot of the SNP density is plotted on the righthand, y-axis showing a single horizontal line for each SNP in the supplied genotype file. A point is plotted next to gene labels to give an indication of gene level importance. The variable to assign this color is also user-specified (**Figure 1**).

**Figure 1:**
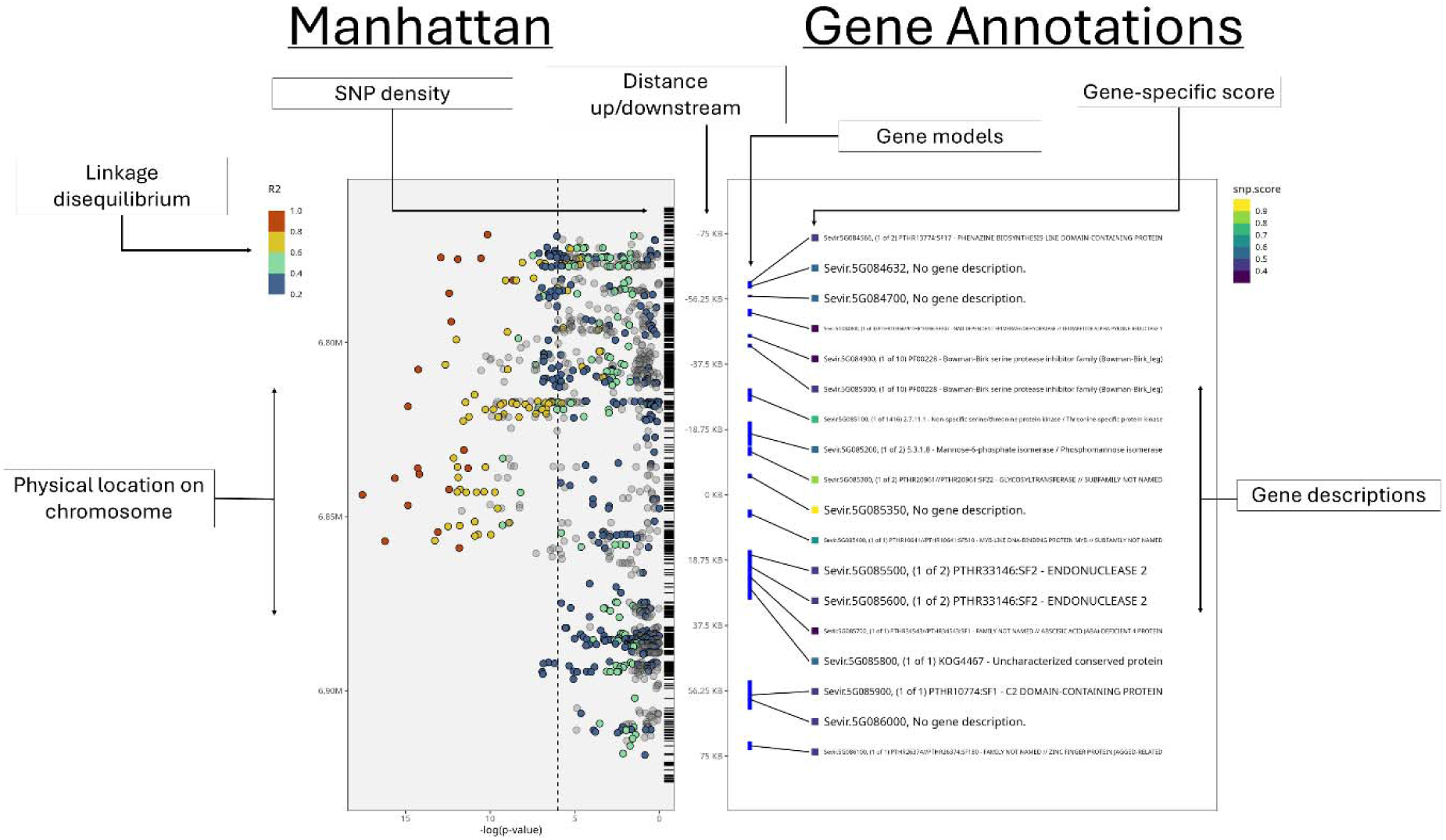
An example of the output of plot_panvar(), the standard visualization for the package. The program plots the Manhattan plot, vertically oriented, and genes in the region side by side. Points in the Manhattan can be colored by snp-specific quantitative measures, by default this is linkage disequilibrium. A rug style plot of snp density in the plotted region is included at the bottom of the Manhattan. Gene locations and annotations are plotted in the right-side panel with points that can be colored by gene-specific quantitative measurements.

### Shiny

A shiny implementation of the pipeline is presented for users that prefer a more interactive experience. Users must go through pre-processing steps up to the generation of the output tables before using the app including formatting GWAS results or optionally re-running GWAS (**Figure 2**). The shiny app will create tables and plot the results (*panvar_shiny()*) (**Figure 3**).

**Figure 2:**
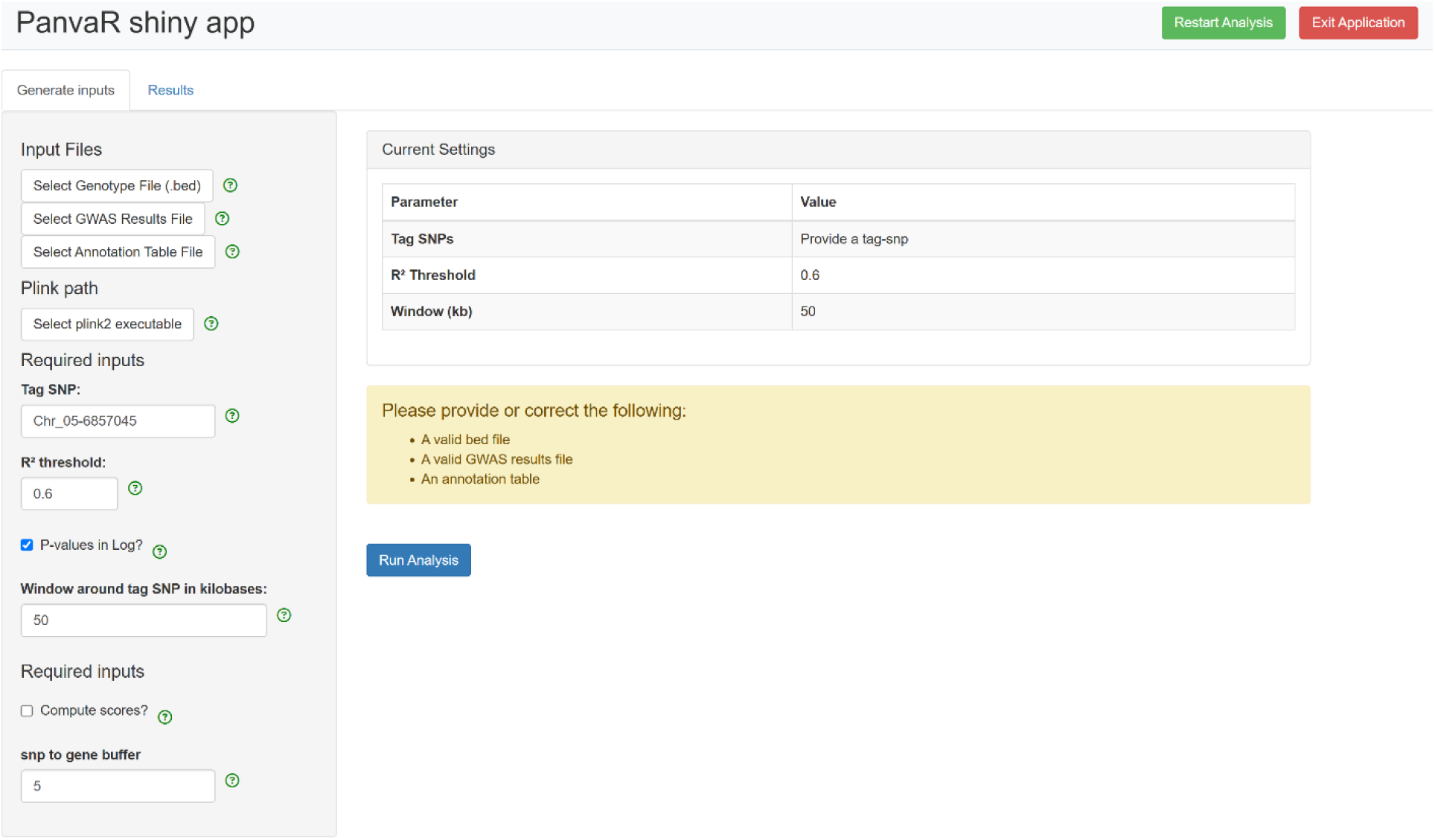
A screenshot of panvaR’s shiny implementation showing the inputs tab. Users input the required files and indicate some options before moving on to generating the outputs. Tooltips can be moused over to provide context for each field.

**Figure 3:**
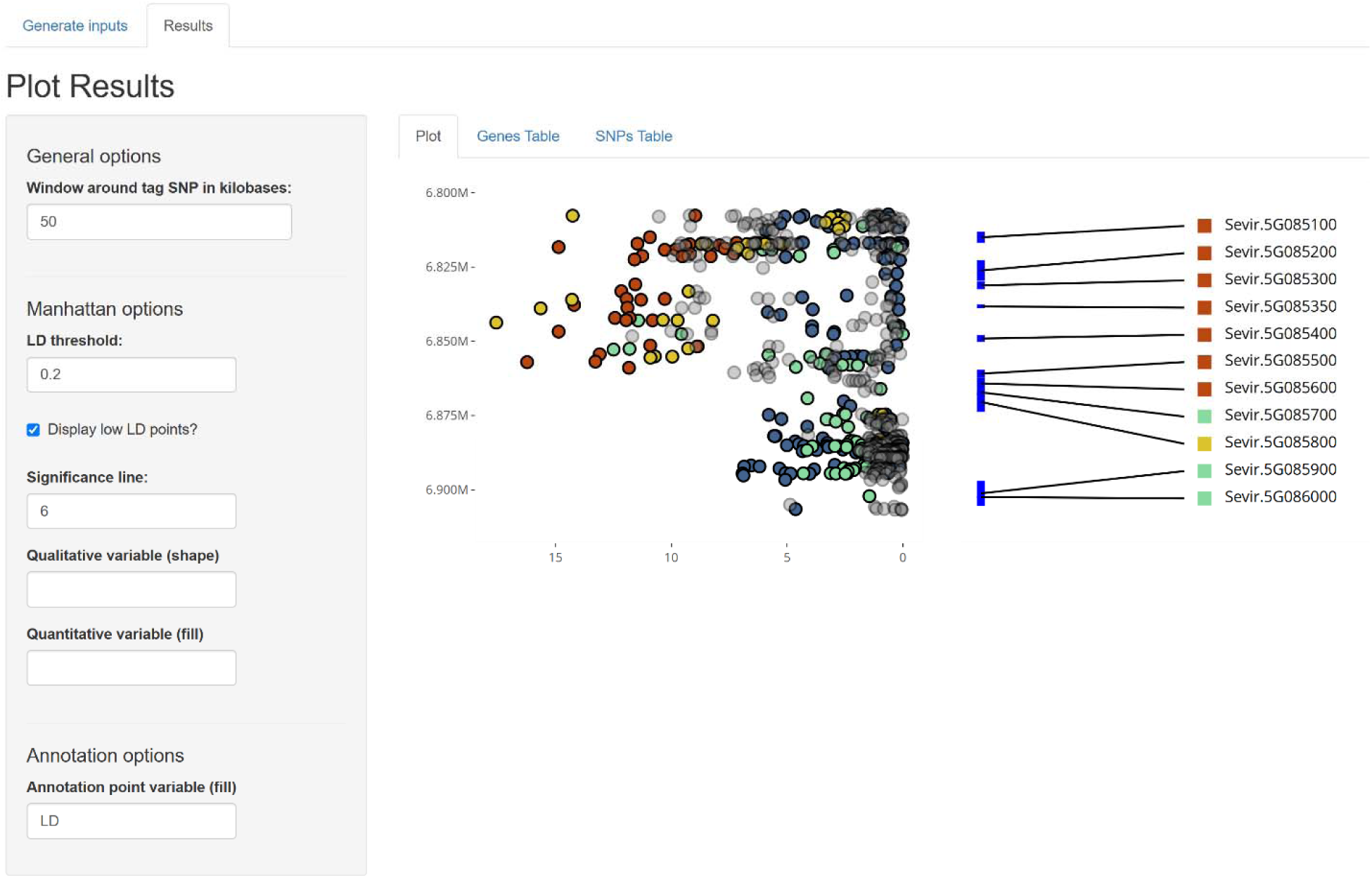
A screenshot of panvaR’s shiny implementation showing the results tab. The results tab displays table and plot output. The plot is interactive and hovering over different parts with the mouse provides more information. Some options for plotting are available for user customization.

## Results

To validate the ability of panvaR to identify previously published genes we present two case studies.

The first case study involved the SvLes1 gene (Sevir.5G085400) identified by Mamidi et al. (2020) (Mamidi et al. 2020). We ran a GWAS and identified the most significant p-value SNP at 6,857,045 base pairs on chromosome 5, which corresponded with the large GWAS peak shown in the paper (**Figure 4A**). We ran panvaR with a 75 kilobase window up and downstream of this SNP (**Figure 4B**). After generating SNP scores using the default distance, p-value and LD to rank SNPs and aggregating SNPs to the gene level, the SvLes1 gene was ranked second of 19 genes in the window. Mamidi et al. took a similar approach in that they identified protein-disrupting polymorphisms using snpEff. However, they did not consider the added information of linkage disequilibrium and instead relied on physical proximity alone to define the size of their GWAS peak. These contrasts highlight panvaR’s utility as a data integration tool capable of both fitting into and extending an already established framework for finemapping.

**Figure 4:**
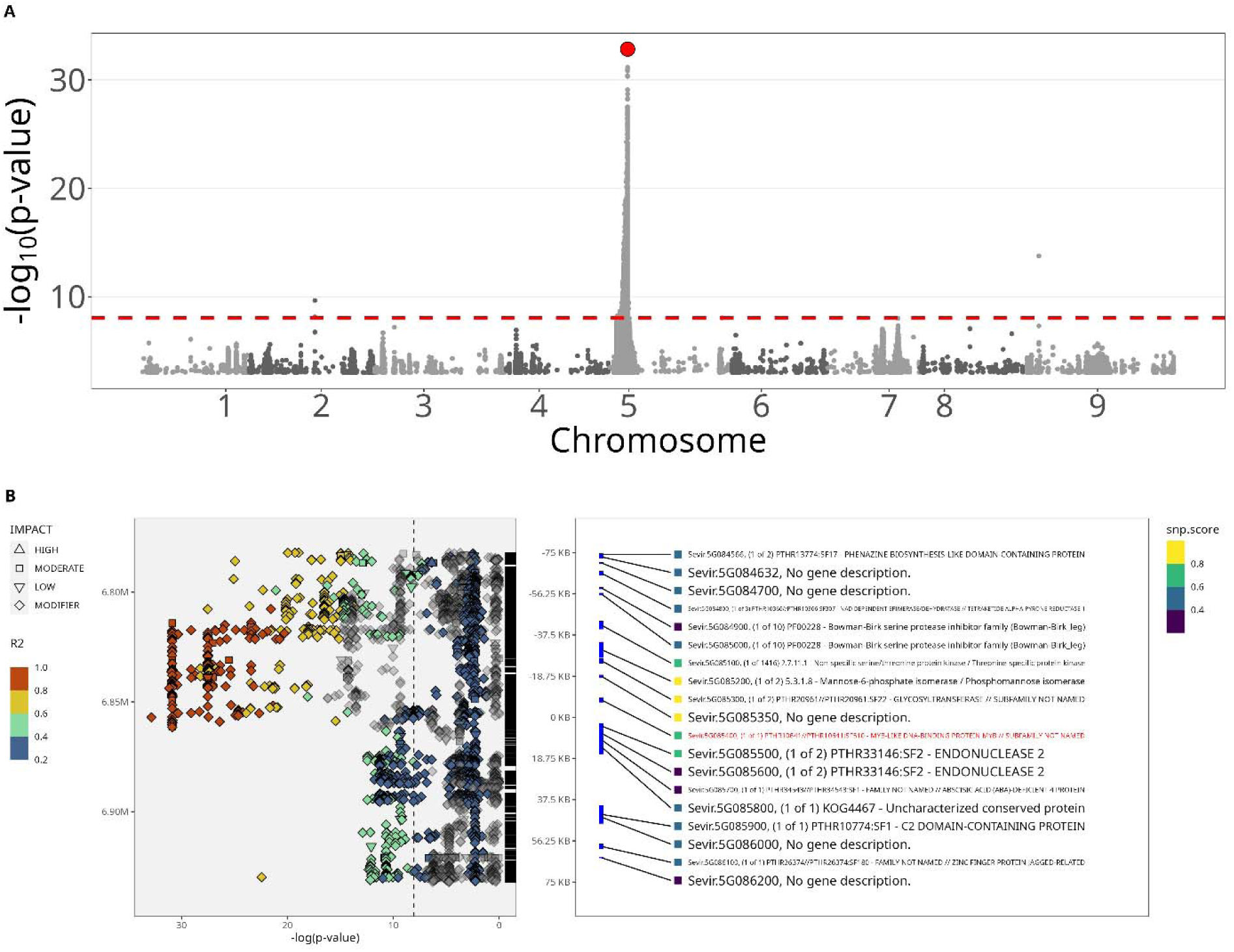
GWAS results and panvaR visualization of identified peak in Setaria case study. A) Manhattan plot of shattering in setaria with the top p-value SNP plotted in red. B) Output from panvaR using the top SNP from GWAS. Gene identified by the paper is highlighted in red.

The second case study involved Brenton et al. (2020) (Brenton et al. 2020), which identified Sobic.004G301500, a vacuolar iron transporter, as a key gene influencing nonstructural carbohydrate levels in Sorghum bicolor. We used a more current version of the sorghum genome as input into the algorithm than was used in Brenton et al. We first uplifted version 3 coordinates to version 5, then ran a GWAS using panvaR. We identified the most significant SNP on chromosome 4 corresponding to the one found in the paper (**Figure 5A**). This SNP, at a physical position of 66,529,675 base pairs, was used as the tag SNP in panvaR with a window of 75 kilobases (**Figure 5B**). For this set of SNPs, we generated variant effect scores from a plant DNA language model, Plant Caduceus (Zhai et al. 2025). These scores assign a numeric value to each SNP’s predicted conservation.

**Figure 5:**
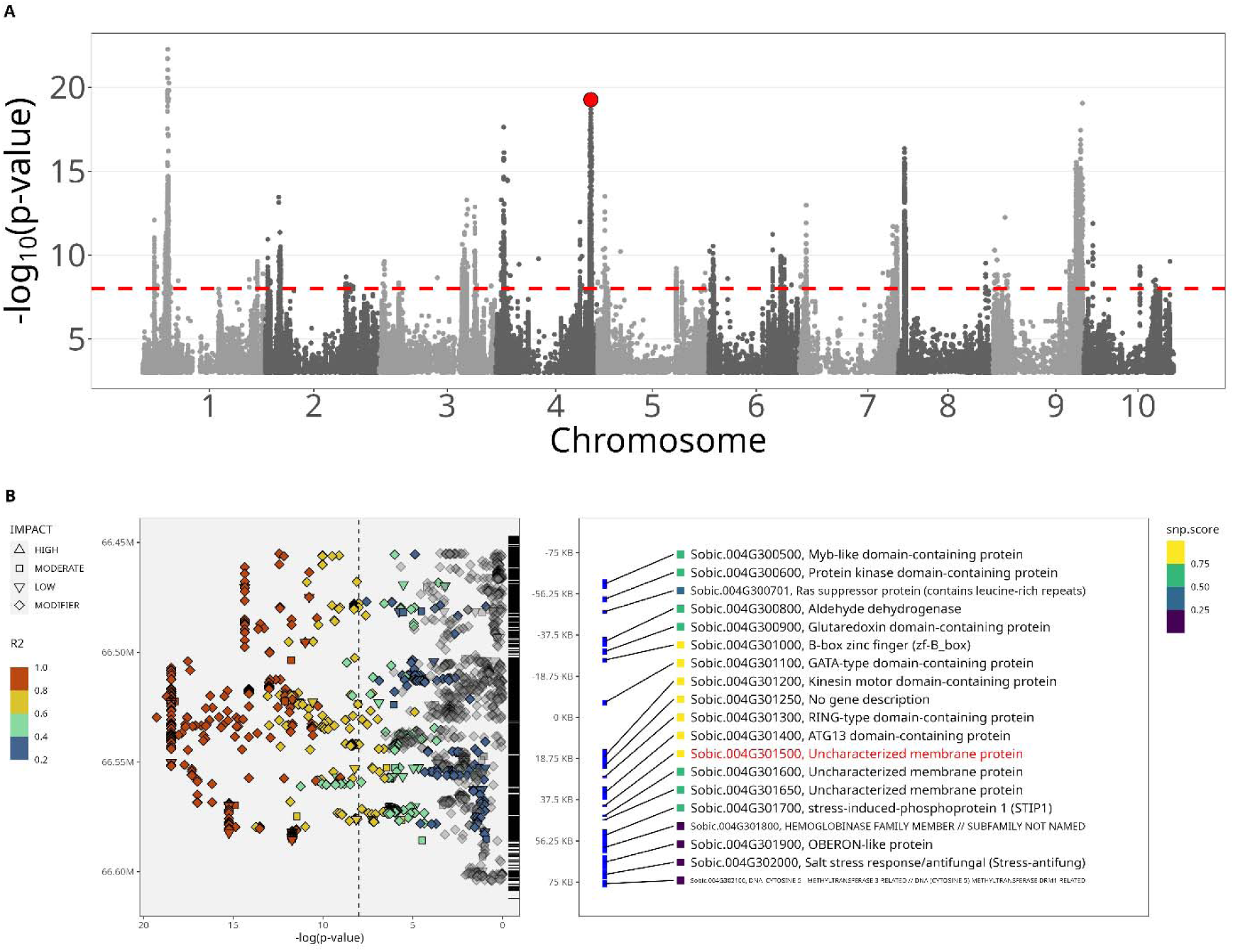
GWAS results and panvaR visualization of identified peak in Sorghum case study. A) GWAS of nonstructural carbohydrate levels in sorghum. The most significant snp in the peak on chromosome 4 that Brenton et al identified is plotted in red. B) Results of panvaR run on the SNP from A. The gene identified in Brenton et al is highlighted in red.

PanvaR can aggregate snp-level metrics like these variant scores to the gene level by choosing the maximum value within an interval of the start and end of a gene. Using these gene-level variant scores, the gene identified in the paper ranked 5th out 19 genes in the window. The other 18 top-ranked genes were identified in Brenton et al. as possible candidates (**supplemental table 1**). The addition of a novel, quantitative measure of SNP importance adds a granularity to the ranking that was absent in the original analysis. This displays panvaR’s utility as an out-of-the-box tool to perform commonplace analyses and its flexibility to include novel and arbitrary measures of SNP importance.

## Discussion

As our two case studies indicate, panvaR speeds up fine-mapping a GWAS peak, producing an easily reviewable list for the user and high-quality visualizations. By synthesizing popular tools, panvaR is able to streamline the display of common fine-mapping data streams like gene locations and linkage disequilibrium information. With its flexible nature in accepting inputs from any species and its extendability in accepting arbitrary measures of SNP importance, panvaR serves as a platform for easily navigating the process of linking GWAS to causal genes.

Ease of use is a prime motivation for panvaR’s design but its framework also has other benefits. Multi-locus models have become increasingly popular in plant GWAS as they effectively control false positives while simultaneously limiting false negatives (Liu et al. 2016). This balance has proven an effective tool for finding strong trait associations (Ward et al. 2019), but information about the significance of phenotypic association is collapsed into a single SNP in subsequent fine-mapping. By running a more permissive, single-locus model, panvaR fills in the information that was collapsed during the model-fitting process, giving more insight into the structure of a GWAS peak than using only p-values from the multi-locus model. Another context in which this filling in of p-values might be useful is if the GWAS was run on a reduced SNP set to conserve computational resources. PanvaR makes it easier to move from a reduced SNP set to focus on all of the pan genomic data for a single locus.

PanvaR is intended as a data integration tool and its gene prioritization system relies on a weighted average of user-defined factors. As such, the list output by the tool should still be inspected manually for best results. Even so, this coarse approximation performs well in our two case studies, where we successfully identified two known genes within a small number of previously top-ranked genes. PanvaR is intentionally designed to be flexible to whichever tools the user wants to incorporate into the scoring system meaning it can easily accommodate novel measures of both SNP and gene importance. With the speed that DNA language models are evolving, it is likely that more performant importance metrics will be released over the coming years (Zhai et al. 2025).

PanvaR is designed to visualize linkage disequilibrium of one or a small number of tag SNPs to all other SNPs in a region. However, more complicated LD structures can exist within a GWAS-identified region which are not displayed using this approach. Visualizing a full pair-wise LD matrix is often useful as it can give insight into more complex inheritance patterns present in the region of interest (Larsson, Lipka, and Buckler 2013). Furthermore, this focus on a single lead SNP can lead to spurious identification of causal genes caused by the phenomenon of “synthetic association” (Platt, Vilhjálmsson, and Nordborg 2010). This can occur either when a rare causal variant is linked to a more common variant or there are multiple genes in the region controlling the trait of interest (Dickson et al. 2010) (Sasaki et al. 2021). These more complex forms of linkage are not detectable within the panvaR framework. That being said, a future direction of the tool will look to include visualizations of the full LD structure in the region and more comprehensive haplotype analysis which should give more insight to the user in these instances.

As a data integration tool, Panvar does not systematically combine data in a statistical way as Bayesian (Chen et al. 2016) and frequentist (Cho et al. 2009) statistical fine-mapping tools do. These tools have been used extensively in human genetics reports of their efficacy in plants has not been comprehensively validated (Wu et al. 2022). Some hurdles to the use of these statistical fine mapping approaches include computation time (Chen et al. 2016), inconsistency of results between different models and a tendency towards false negatives depending on the model (Schaid, Chen, and Larson 2018). Despite these hurdles, these statistical fine-mapping could be integrated into the framework of Panvar as additional decision making criteria in causal gene identification as they become more developed for use in plants.

## Conclusions

PanvaR synthesizes data types commonly used in fine-mapping GWAS peaks to speed up this step in the gene discovery pipeline. Information afforded by pangenomic resources can easily be mined around previously identified peaks, further adding to panvaR’s utility. Going forward, panvaR should be a worthy addition to the quantitative genetic researchers’ toolbox, enabling novel insights into genes and their functions.

## Bibliography

Brenton, Zachary W., Brendon T. Juengst, Elizabeth A. Cooper, Matthew T. Myers, Kathleen E. Jordan, Savanah M. Dale, Jeffrey C. Glaubitz, et al. 2020. “Species-Specific Duplication Event Associated with Elevated Levels of Nonstructural Carbohydrates in Sorghum Bicolor.” G3 (Bethesda, Md.) 10 (5): 1511–20.

Chang, Christopher C., Carson C. Chow, Laurent Cam Tellier, Shashaank Vattikuti, Shaun M. Purcell, and James J. Lee. 2015. “Second-Generation PLINK: Rising to the Challenge of Larger and Richer Datasets.” GigaScience 4 (1): 7.

Chen, Wenan, Shannon K. McDonnell, Stephen N. Thibodeau, Lori S. Tillmans, and Daniel J. Schaid. 2016. “Incorporating Functional Annotations for Fine-Mapping Causal Variants in a Bayesian Framework Using Summary Statistics.” Genetics 204 (3): 933–58.

Cho, Seoae, Haseong Kim, Sohee Oh, Kyunga Kim, and Taesung Park. 2009. “Elastic-Net Regularization Approaches for Genome-Wide Association Studies of Rheumatoid Arthritis.” BMC Proceedings 3 Suppl 7 (S7): S25.

Collard, Bertrand C. Y., and David J. Mackill. 2008. “Marker-Assisted Selection: An Approach for Precision Plant Breeding in the Twenty-First Century.” Philosophical Transactions of the Royal Society of London. Series B, Biological Sciences 363 (1491): 557–72.

Della Coletta, Rafael, Yinjie Qiu, Shujun Ou, Matthew B. Hufford, and Candice N. Hirsch. 2021. “How the Pan-Genome Is Changing Crop Genomics and Improvement.” Genome Biology 22 (1): 3.

Dickson, Samuel P., Kai Wang, Ian Krantz, Hakon Hakonarson, and David B. Goldstein. 2010. “Rare Variants Create Synthetic Genome-Wide Associations.” PLoS Biology 8 (1): e1000294.

Guo, Dongling, Yan Li, Hengyun Lu, Yan Zhao, Nori Kurata, Xinghua Wei, Ahong Wang, et al. 2025. “A Pangenome Reference of Wild and Cultivated Rice.” Nature 642 (8068): 662–71.

Hufford, Matthew B., Arun S. Seetharam, Margaret R. Woodhouse, Kapeel M. Chougule, Shujun Ou, Jianing Liu, William A. Ricci, et al. 2021. “De Novo Assembly, Annotation, and Comparative Analysis of 26 Diverse Maize Genomes.” Science (New York, N.Y.) 373 (6555): 655–62.

Kang, Minghui, Haolin Wu, Huanhuan Liu, Wenyu Liu, Mingjia Zhu, Yu Han, Wei Liu, et al. 2023. “The Pan-Genome and Local Adaptation of Arabidopsis Thaliana.” Nature Communications 14 (1): 6259.

Larsson, Sara J., Alexander E. Lipka, and Edward S. Buckler. 2013. “Lessons from Dwarf8 on the Strengths and Weaknesses of Structured Association Mapping.” PLoS Genetics 9 (2): e1003246.

Liu, Xiaolei, Meng Huang, Bin Fan, Edward S. Buckler, and Zhiwu Zhang. 2016. “Iterative Usage of Fixed and Random Effect Models for Powerful and Efficient Genome-Wide Association Studies.” PLoS Genetics 12 (2): e1005767.

Mamidi, Sujan, Adam Healey, Pu Huang, Jane Grimwood, Jerry Jenkins, Kerrie Barry, Avinash Sreedasyam, et al. 2020. “A Genome Resource for Green Millet Setaria Viridis Enables Discovery of Agronomically Valuable Loci.” Nature Biotechnology 38 (10): 1203–10.

Morris, Geoffrey P., Avril M. Harder, Adam L. Healey, Chloee M. McLaughlin, Joanna L. Rifkin, Clara Cruet-Burgos, Jerry W. Jenkins, et al. 2026. “A Sorghum Pangenome Reference Improves Global Crop Trait Discovery.” Nature, March. 10.1038/s41586-026-10229-9.

Platt, Alexander, Bjarni J. Vilhjálmsson, and Magnus Nordborg. 2010. “Conditions under Which Genome-Wide Association Studies Will Be Positively Misleading.” Genetics 186 (3): 1045–52.

Sasaki, Eriko, Thomas Köcher, Danièle L. Filiault, and Magnus Nordborg. 2021. “Revisiting a GWAS Peak in Arabidopsis Thaliana Reveals Possible Confounding by Genetic Heterogeneity.” Heredity 127 (3): 245–52.

Schaid, Daniel J., Wenan Chen, and Nicholas B. Larson. 2018. “From Genome-Wide Associations to Candidate Causal Variants by Statistical Fine-Mapping.” Nature Reviews. Genetics 19 (8): 491–504.

Shi, Junpeng, Zhixi Tian, Jinsheng Lai, and Xuehui Huang. 2023. “Plant Pan-Genomics and Its Applications.” Molecular Plant 16 (1): 168–86.

Tao, Yongfu, Hong Luo, Jiabao Xu, Alan Cruickshank, Xianrong Zhao, Fei Teng, Adrian Hathorn, et al. 2021. “Extensive Variation within the Pan-Genome of Cultivated and Wild Sorghum.” Nature Plants 7 (6): 766–73.

Tibbs Cortes, Laura, Zhiwu Zhang, and Jianming Yu. 2021. “Status and Prospects of Genome-Wide Association Studies in Plants.” The Plant Genome 14 (1): e20077.

Voytas, Daniel F., and Caixia Gao. 2014. “Precision Genome Engineering and Agriculture: Opportunities and Regulatory Challenges.” PLoS Biology 12 (6): e1001877.

Ward, Brian P., Gina Brown-Guedira, Frederic L. Kolb, David A. Van Sanford, Priyanka Tyagi, Clay H. Sneller, and Carl A. Griffey. 2019. “Genome-Wide Association Studies for Yield-Related Traits in Soft Red Winter Wheat Grown in Virginia.” PloS One 14 (2): e0208217.

Whitt, Lauren, Elizabeth H. Mahood, Greg Ziegler, Collin Luebbert, Jason D. Gillman, Gareth J. Norton, Adam H. Price, David E. Salt, Brian P. Dilkes, and Ivan Baxter. 2023. “A Comparative Approach for Selecting Orthologous Candidate Genes in Genome-Wide Association Studies across Multiple Species.” bioRxiv. 10.1101/2023.10.05.561051.

Wu, Xing, Wei Jiang, Christopher Fragoso, Jing Huang, Geyu Zhou, Hongyu Zhao, and Stephen Dellaporta. 2022. “Prioritized Candidate Causal Haplotype Blocks in Plant Genome-Wide Association Studies.” PLoS Genetics 18 (10): e1010437.

Yin, Lilin, Haohao Zhang, Zhenshuang Tang, Jingya Xu, Dong Yin, Zhiwu Zhang, Xiaohui Yuan, et al. 2021. “RMVP: A Memory-Efficient, Visualization-Enhanced, and Parallel-Accelerated Tool for Genome-Wide Association Study.” Genomics, Proteomics & Bioinformatics 19 (4): 619–28.

Zhai, Jingjing, Aaron Gokaslan, Yair Schiff, Ana Berthel, Zong-Yan Liu, Wei-Yun Lai, Zachary R. Miller, et al. 2025. “Cross-Species Modeling of Plant Genomes at Single-Nucleotide Resolution Using a Pretrained DNA Language Model.” Proceedings of the National Academy of Sciences of the United States of America 122 (24): e2421738122.

