## Supplementary figures and images for "PanvaR: An R package for fine-mapping and visualizing results from genome-wide association studies"

### Supplemental Figure 1

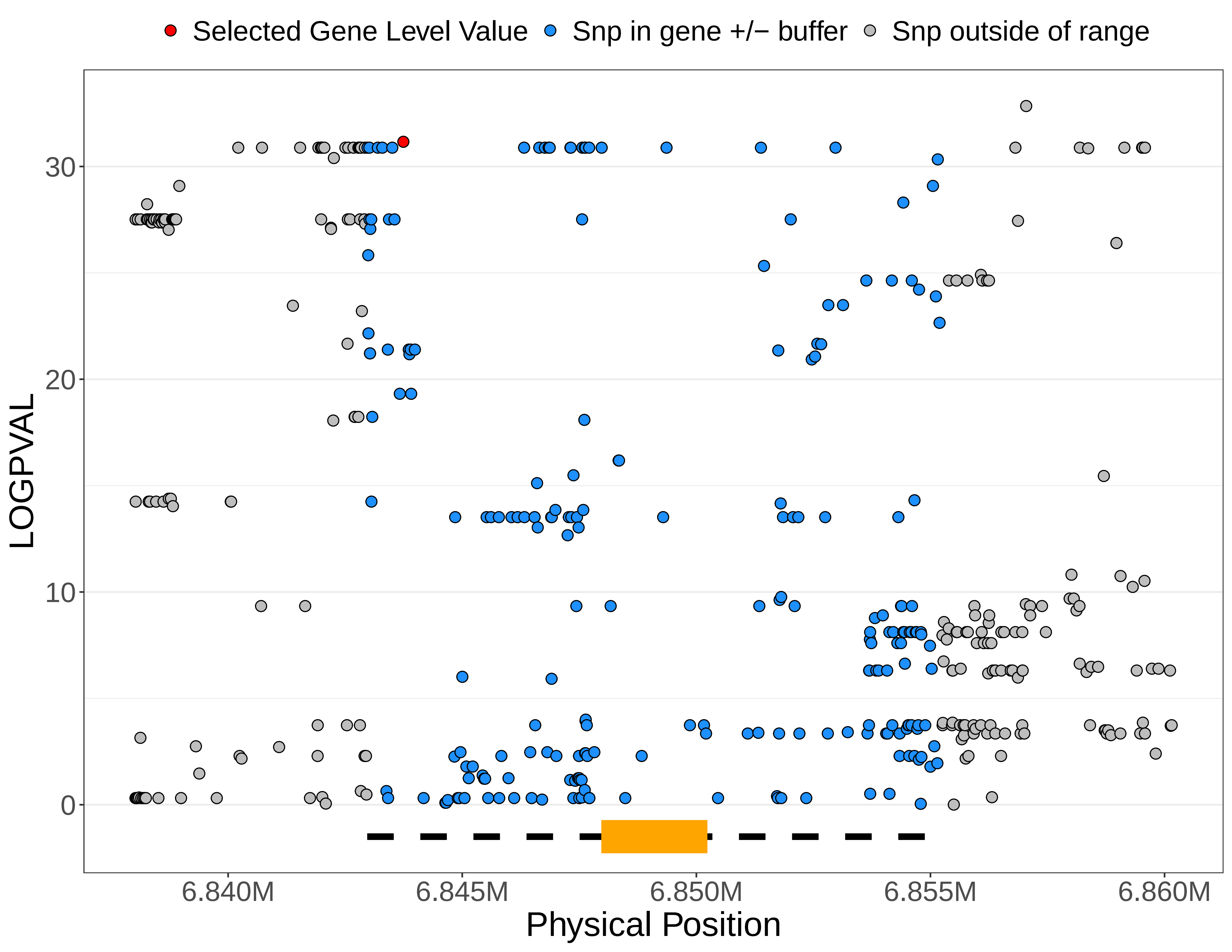
